# ShrinkCRISPR : A flexible method for differential fitness analysis of CRISPR-Cas9 screen data

**DOI:** 10.1101/2022.03.31.486584

**Authors:** Renaud L.M. Tissier, Janne J.M. van Schie, Rob M.F. Wolthuis, job de Lange, Renée X. de Menezes

## Abstract

CRISPR screens provide large-scale assessment of cellular gene functions. Pooled libraries typically consist of several single guide RNAs (sgRNAs) per gene, for a large number of genes, which are transduced in such a way that every cell receives at most one sgRNA, resulting in the disruption of a single gene in that cell. This approach is often used to investigate effects on cellular fitness, by measuring sgRNA abundance at different time points. Comparing gene knockout effects between different cell populations is challenging due to variable cell-type specific parameters and between replicates variation. Failure to take those into account can lead to inflated or false discoveries.

We propose a new, flexible approach called ShrinkCRISPR that can take into account multiple sources of variation. Impact on cellular fitness between conditions is inferred by using a mixed-effects model, which allows to test for gene-knockout effects while taking into account sgRNA-specific variation. Estimates are obtained using an empirical Bayesian approach. ShrinkCRISPR can be applied to a variety of experimental designs, including multiple factors. In simulation studies, we compared ShrinkCRISPR results with those of drugZ and MAGeCK, common methods used to detect differential effect on cell fitness. ShrinkCRISPR yielded as many true discoveries as drugZ using a paired screen design, and outperformed both drugZ and MAGeCK for an independent screen design. Although conservative, ShrinkCRISPR was the only approach that kept false discoveries under control at the desired level, for both designs. Using data from several publicly available screens, we showed that ShrinkCRISPR can take data for several time points into account simultaneously, helping to detect early and late differential effects.

ShrinkCRISPR is a robust and flexible approach, able to incorporate different sources of variations and to test for differential effect on cell fitness at the gene level. These improve power to find effects on cell fitness, while keeping multiple testing under the correct control level and helping to improve reproducibility. ShrinkCrispr can be applied to different study designs and incorporate multiple time points, making it a complete and reliable tool to analyze CRISPR screen data.

## Introduction

The study of effects of genetic perturbation is fundamental to elucidate gene function. In addition, the identification of genes which knockout leads to cell death, either in combination with another genetic change (say a mutation of another gene, leading to ‘synthetic lethality’) or in combination with a certain drug, may lead to more efficient cancer treatments. Genome-scale screening methods, in which thousands of genes are individually targeted in a single experiment, are often at the start of such investigations. Major challenges of such approaches have included undesired targeting of aspecific sites (“off-target” effects) and variable gene inactivation efficiencies. The adaptation of clustered regularly interspaced short palindromic repeats (CRISPR) technology to mammalian cells led to the development of improved gene knockout screens, with higher efficiency and lower off-target effects [7, 8].

Per cell one gene is knocked out and single guide RNAs (sgRNAs) of these cells are sequenced. By comparing sgRNA abundance between different conditions, the effect of specific knockouts on cell fitness can be investigated.

This method can be applied to study the impact of gene knockout on cell lines of different origins, isogenic cell lines (identical cell lines in which only the status of one gene is different) or a cell line with and without treatment. However, the comparison of gene knockout effects is challenging due to differences in abundance of specific sgRNAs at the start of the experiment. These may be due, for example, to variations in the library composition, efficiency of transduction of sgRNAs into cells and selection, growth rate of transduced cells, premature or incomplete *Cas9* activity and between-replicate variation. Data analysis methods ideally should take care of all these issues, to ensure reliability and reproducibility of identified effects.

CRISPR screen data involve additional aspects that need to be taken into account: (i) a large number of variables (sgRNAs) and a relatively small number of replicates, also typical for other omics data; (ii) the data are generated by DNA sequencing, and thus consist of counts displaying over-dispersion, aspects that classic statistical methods do not account for; (iii) the effect of a gene knockout is evaluated by several sgRNAs per gene, which need to be aggregated to reach a single conclusion about that gene; (iv) the experiment often involves two sequencinq runs, one at baseline measuring the starting abundance of each sgRNA, and a paired replicate at a later time point, typically after a chosen number of cell doublings. Furthermore, data related to multiple cell lines exhibit both technical and biological variability, which must be accounted for separate and differently in the data analysis. Indeed, the effect of condition such as different treatments may be represented as fixed in the model, as conclusions are to be drawn for the chosen conditions, whilst the variation between sgRNAs can better be assumed to be random – the sgRNA effect then represents that of multiple, similar sgRNAs targeting the same gene.

Currently available analysis methods only account for some of these issues. For example, two often used methods, MAGeCK [11] and drugZ [3], ignore the variability between initial replicates. Since the data are produced using deep sequencing, some authors have used data analysis methods designed for (RNA) sequencing, such as edgeR [4, 13, 17] and DESeq2 [12]. While these can take care of over-dispersion, neither can estimate both the fixed effect of condition, as well as the random effect of sgRNAs. Furthermore, as these perform one test per sgRNA, they typically require a much heavier multiple testing correction than if testing was performed per gene.

To tackle these challenges, we propose an analysis method that, by means of a mixed-effects regression-based empirical-Bayes framework, can efficiently detect genes with differential impact on cell fitness. It first transforms the count data into fold changes relative to the starting sgRNA abundance at *T* = 0 for each biological sample, then normalizes the data using the rscreenorm [1] method. It then uses an empirical-Bayes regression model, including a fixed effect for condition and a random effect for the sgRNAs, to find gene-specific effects. This leads to a test per gene by taking all sgRNAs targeting that gene together into account. Our method makes use of ShrinkBayes [19], which fits empirical-Bayes regression models simultaneously for many features using INLA [18], enabling the use of mixed effect models in analyses of high-dimensional data. We call our method *ShrinkCRISPR*.

This manuscript is organized as follows. Section 2 presents the method, section 3 uses a simulation model to compare methods’ performances and section 4 illustrates the performance of our pipeline on experimental data. We conclude with a brief summary and discussion in section 5.

## Methods

### Experimental designs

For completeness, we give here a short overview of the steps involved in CRISPR-Cas9 screening. For each replicate, viral particles expressing single guide RNAs (sgRNAs) are transduced into cells, and each sgRNA leads the Cas9 endonuclease to cut a specific target, resulting in a specific gene knockout [8]. Transduction efficiency can vary, so abundance of each sgRNA is obtained per sample both at an initial time point, which we will refer to as *T* = 0, as well as at a later time point, typically after a chosen number of cell doublings. Differences in sgRNA abundance between the investigated experimental conditions reflect a differential effect on cell fitness of the specific knockout.

Two experimental designs are commonly used when studying differential effect on cell fitness between cell lines. For the assessment of drug sensitivity, one batch of cells is transduced and sequenced (*T* = 0) after which the cell sample is split, and the resulting parts are cultured separately, one with and the other without the drug of interest. This experimental design yields data that can be analysed per pair of replicates disregarding initial sgRNA abundance at *T* = 0, as this is the same for both. Data analysis involves comparing sgRNA abundance between the treated and untreated replicates (Figure 1-A). We will refer to this drug sensitizing experimental design as the “paired design”.

**Figure 1.**
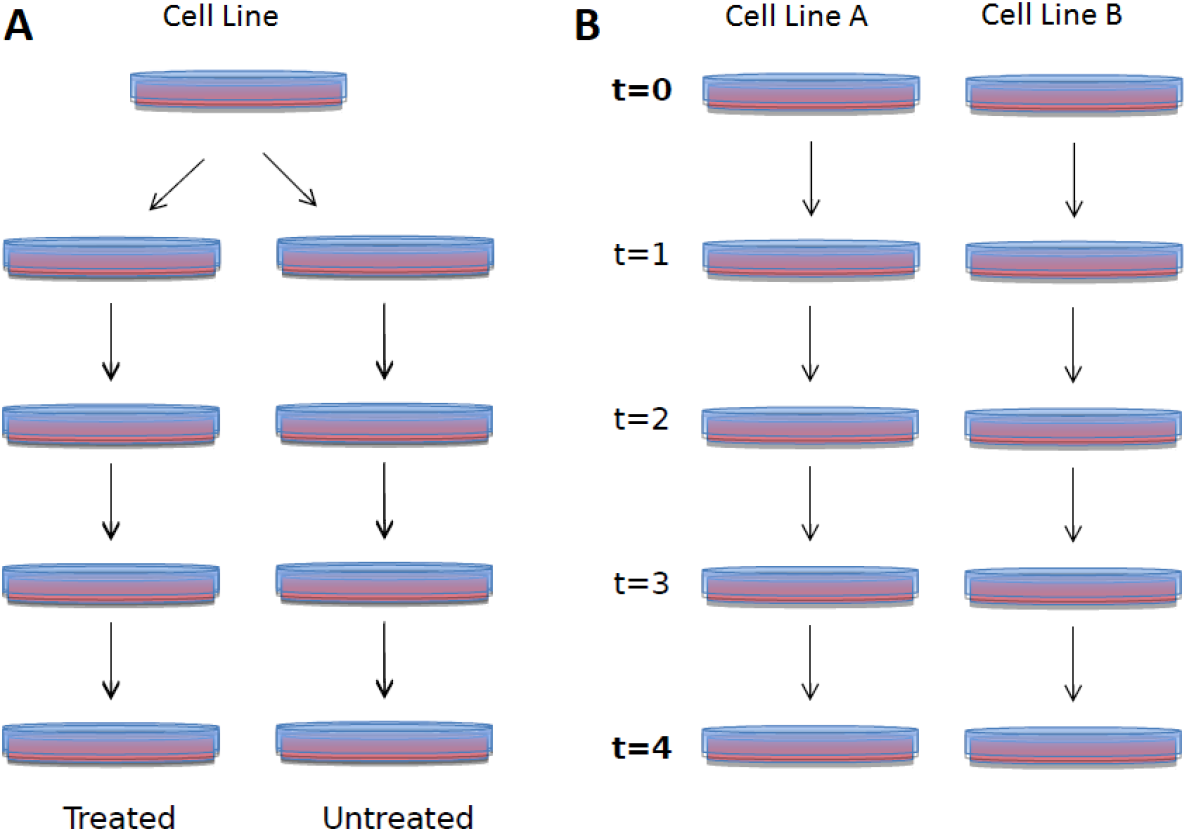
Two commonly used experimental designs. A: a drug sensitizing study compares the effect of gene knockout on paired replicates derived from the same initial sample screened at *T* = 0, which is then split and each part is cultured under a different condition (typically treatment or not). B: the comparison of two cell lines typically involves different sgRNA abundances at *T* = 0.

For comparing two different cell lines, initial sgRNA abundance at *T* = 0 is measured per cell line. Then sgRNA abundances are compared between cell lines A and B at a later time point (Figure 1-B). We will refer to this design as the the “independent design”. The two cell lines here involved can for example be derived from the same parental cell line, with and without a mutation.

### Taking *T* = 0 into account and normalization

When using the independent design, sgRNA abundance can only be compared fairly between cell lines if its initial abundance is taken into account. To do this, we transform the count data into fold changes using the formula:

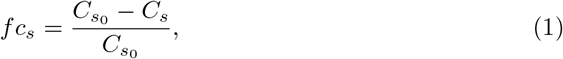

for a given sgRNA *s*, where *C*_*s*_0__, *C_s_* represent the cell count at *T* = 0 and at the end point of the screen, respectively, and *fc_s_* is the resulting fold change. Indeed, *fc_s_* is the proportion of cells containing sgRNA *s* lost relative to their initial abundance. It can take values from –∞ to 1, where 1 represents complete lethality (all cells with sgRNA *s* are lost between *T*_0_ and the end point, so *C_s_* = 0), 0 represents no effect on cell viability (i.e. *C_s_* = *C*_*s*_0__), and negative values indicate proliferation of cells containing the sgRNA s (*C_s_* > *C*_*s*_0__).

The distribution of fold changes can vary between replicates, between conditions and between cell lines. The use of normalization procedures such as rscreenorm [1] is especially important when these distributions vary considerably. Rscreenorm makes use of assay controls (both negative and positive controls) and of quantile normalization to make measurements across replicates and cell lines more comparable and reflecting the functional effect. With rscreenorm we generate *lethality scores* per sgRNA, with values around 0 for sgRNAs yielding a phenotype similar to that of negative controls, and values around 1 for sgRNAs similar to positive controls. In other words, lethality scores around 0 represent the same cell fitness as for negative controls, whilst lethality scores around 1 represent as little cell fitness as for positive controls. Quantile normalization is only required in case of large differences in fitness score distributions of the different samples. Lethality scores *l_s_* are derived from the fold changes via the formula:

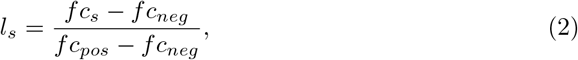

where *fc_neg_* is the median of the sgRNAs’ fold changes in the negative control genes and *fc_pos_* is the median of the sgRNAs’ fold changes in the positive control genes.

### An empirical-Bayes regression model

By testing possible differences between fold changes of the different cell lines or conditions, we can detect differential effect on cell fitness. To do so, we define a multivariate statistical model. For given gene *g*, replicate *r* and condition *c*, the following model is used:

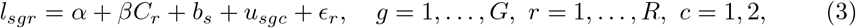

where *s* ∈ {1,…, *S_g_*} indicates the sgRNA targeting gene *g*, and *S_g_* is the total number of sgRNAs targeting gene *g*. In addition, *l_sgr_* is a vector of length equal to *S_g_*, *α* is the vector of average lethal effects of sgRNAs in the reference cell line, *β* represents the difference in lethality of the gene knockout between the two conditions across sgRNAs targeting gene *g*, *C_r_* represents the condition for replicate *r*, *b_s_* is a random effect accounting for variation between sgRNAs, and *u_sgc_* is a random effect representing a possible interaction between sgRNA *s* and condition *c*, allowing for condition-specific sgRNA effects. The error term *ϵ_r_* of the model is assumed to follow a normal distribution with mean 0. We can then test the null hypothesis *H*_0_: *β* = 0 of no differential lethality of any of the sgRNAs s between the conditions, against the alternative hypothesis *H_a_*: *β* ≠ 0 of differential lethality. If an independent design is used, the condition represents a specific cell line, and the remaining cell line is used as reference. If a paired design is used, the condition represents a treated sample, and the untreated sample is used as reference.

This flexible model tries to capture multiple sources of variation at the gene level, namely across replicates (by means of *C_r_*) and sgRNAs (via *b_s_* and *u_sgc_*), yielding a single test per gene for difference in cell fitness between conditions. The model can be extended to incorporate longitudinal data.

Experiments involving CRISPR screens often involve a low number of replicates over just two conditions, while measuring the abundance of tens of thousands of sgRNAs at the same time. Model fitting per sgRNA may therefore lead to unreliable estimates. To counter this, we fit the model using ShrinkBayes [19], an empirical-Bayes approach which uses shrinkage to yield parameter estimates. This produces estimates of fixed as well as of random effects in the model by means of efficient, deterministic numerical approximations using INLA [18]. Because our proposed method involves shrinkage, we call it “ShrinkCRISPR”. An R package is currently under construction for the use of ShrinkCRISPR and all codes used in this paper are available on the github page https://github.com/RenTissier.

### Per gene testing

ShrinkCRISPR enables us to test for differential effects on cell fitness directly at the gene level. This is equivalent to testing *H*_0_: *β* = 0 in equation 3. As such, the test takes into account existing variation between sgRNAs and between replicates, as well as a possible interaction between sgRNAs and conditions. By taking all sgRNAs into account and modelling the separate sources of variation, this yields robust effect estimates which tend to be more reproducible.

### Multiple testing correction

For each gene, a Bayes factor is obtained by fitting the model under the null and the alternative hypotheses and computing the ratio between the two posterior marginal likelihoods obtained. From this list of Bayes factors, the local false discovery rate (lfdr, [5]) is computed for each gene. The lfdr has the advantage over the false discovery rate that the Bayes factors involved do not need to be independent.

### Multiple time points

Another strength of ShrinkCRISPR is the possibility to include sgRNA abundance for more than two time points in the model. This can lead to an increase in power, as more measurements are available per replicate, and may allow for a reduction of the number of replicates. Model 3 with multiple time point becomes:

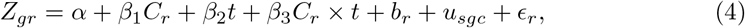

where *β*_1_ represents the average differential lethality effect of the gene knockout between conditions, across sgRNAs and over time. The parameter *β*_2_ can be interpreted as the change in cell population containing sgRNAs targeting gene *g*, and *t* represents the time points used. Another parameter of interest in this model is the interaction effect *β*_3_, representing a possible interaction between time and condition on the fold change of the sgRNAs targeting gene *g*.

## Simulation study

### Simulation setup

We performed a simulation study to evaluate the performance of ShrinkCRISPR, and to compare it to the commonly used methods drugZ [3] and MAGeCK [11]. All methods were compared in terms of true effects detected, as well as false discoveries produced. We simulated data for both paired and independent designs. This allows us not only to compare all three methods, but also to study the impact of variability of initial sgRNA abundance on results, which plays a role when using an independent design, but not when using a paired design. We assume the study involves two conditions (*c* = 1, 2) with *R* replicates in each condition, and *S* sgRNAs studied so that *S* = ∑_*g*_ *S_g_*, with *S_g_* representing the number of sgRNAs per gene as before.

Our simulation setup involved the following steps:

1. Simulation of the mean lethality effect 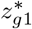 for each gene *g* in the control cell line using a gamma distribution of shape and scale fixed to 1:

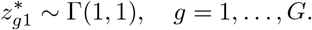 As shown in figure S1 of the supplementary material, such gamma distribution allows for a majority of lethality effects to be close or equal to 0. Assuming that most genes have a behaviour closer from a non-essential gene (gene known to not be impacting cell survival if knocked out) than from an essential gene (gene required for cell survival). The randomly generated lethality effects are subsequently scaled between 0 and 1 by dividing them by the maximum lethality effect obtained. Note that by rescaling the lethality effects, the lethality effect drawn from the gamma distribution are shrank towards zero (as we are divinding by a value higher than 1) subsequently increasing the number of non-essential genes in the control cell line.
2. Computation of the mean *lethality* effect for each gene in condition 2 by using 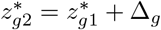, where Δ_*g*_ represents the mean effect difference of all sgRNAs *s* targeting gene *g* (defined later on). We define by 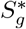 the set of indices *s* of sgRNAs targeting gene *g*. In the context of this simulation study, all sets 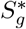 include 4 indices. Furthermore, we represent by *z_sc_* the lethality effect for sgRNA *s* in condition *c*, which is equal to 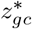 for 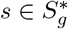. So, {*z_sc_*, *s* = 1,…, } is the expanded set of values of lethality effects over all sgRNAs, corresponding to the original set {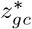, *g* = 1,…, *G*} of generated values for all genes.
3. Simulation of the observed fold changes *fc_src_* for each sgRNA *s*, replicate *r* and condition *c* using a Gaussian distribution:

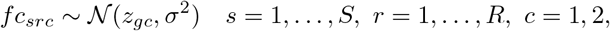

where *σ*^2^ represents the biological variation between replicates of the same condition. The generated values {*fc_src_*, *s* = 1,…, *S*} for each replicate *r* are then organized in vectors, which form the columns of a matrix *L* of dimensions *S* × (2*R*).
4. Simulation of the *S* × (2*R*) matrix *C*_0_ representing the initial counts for each replicate shortly after transduction. The counts for the independent design case are simulated to control the average initial number of counts for each sgRNA as follows:

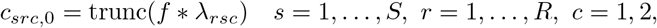

where *f*, i.e, the number of folds represents the average count of cells transduced by each sgRNA, 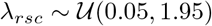 represents the transduction efficiency, and “trunc” is the truncation function. In the paired design case, the equation becomes:

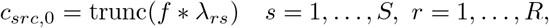
5. Computation of the matrix *M* of gene counts at a later time point using the formula:

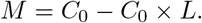

Each dataset contains 2 conditions with *R* = 3 replicates each and *G* = 1000 genes. Simulations are made independently for both paired and independent designs. For each gene, we simulate a fixed number *S_g_* = 4 of different sgRNAs, resulting in a total of *S* = 4000 sgRNAs. In each dataset the first 100 genes (400 sgRNAs) are simulated using the same value Δ_*g*_ as mean effect difference for their sgRNAs. So for each experimental design, the value of Δ_*g*_ is given by:

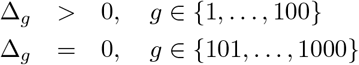

For each experimental design we consider 6 scenarios, each corresponding to a different value of Δ_*g*_ ∈ {0, 0.1, 0.2, 0.3, 0.4, 0.5}. A fixed value of *σ* = 0.1 and *f* = 400 is used in all scenarios. A total of 100 datasets are simulated and analyzed for each scenario. Four hundred positive control sgRNAs (sgRNAs from essential genes) and 400 negative control sgRNAs (sgRNAs from non-essential genes) are also simulated in each dataset. The simulation process is the same for remaining sgRNAs, with the exception that their mean lethality *z*_*g*1_ is fixed and is the same for both conditions. The mean lethality for positive controls is 0.8 and for negative controls is 0.1. A fixed value of *σ* = 0.05 is used for the controls.

ShrinkISO is then applied for each of the 100 simulated datasets, and genes with lfdr < 0.05 are selected as significant.

## Results

### Paired design

Figure 2 shows that the performance obtained by the three approaches is rather different across all simulation scenarios. Indeed, drugZ and MAGeCK yield too many false positive hits even when there are no differential effects, i.e. when Δ_*g*_ =0, with on average 66.9 and 105.2 false hits respectively (left panel of figure 2). These numbers represent 6.7 and 10.5% of false positives, respectively, when in fact 5% were expected. If no effect is simulated, then 100% of the hits are false positives. As Δ_*g*_ increases, drugZ yields progressively less false positives, but that is not the case for MAGeCK. In contrast, ShrinkCRISPR does not yield any false positives across all different scenarios. This suggests it is conservative, in this simulation setup.

**Figure 2.**
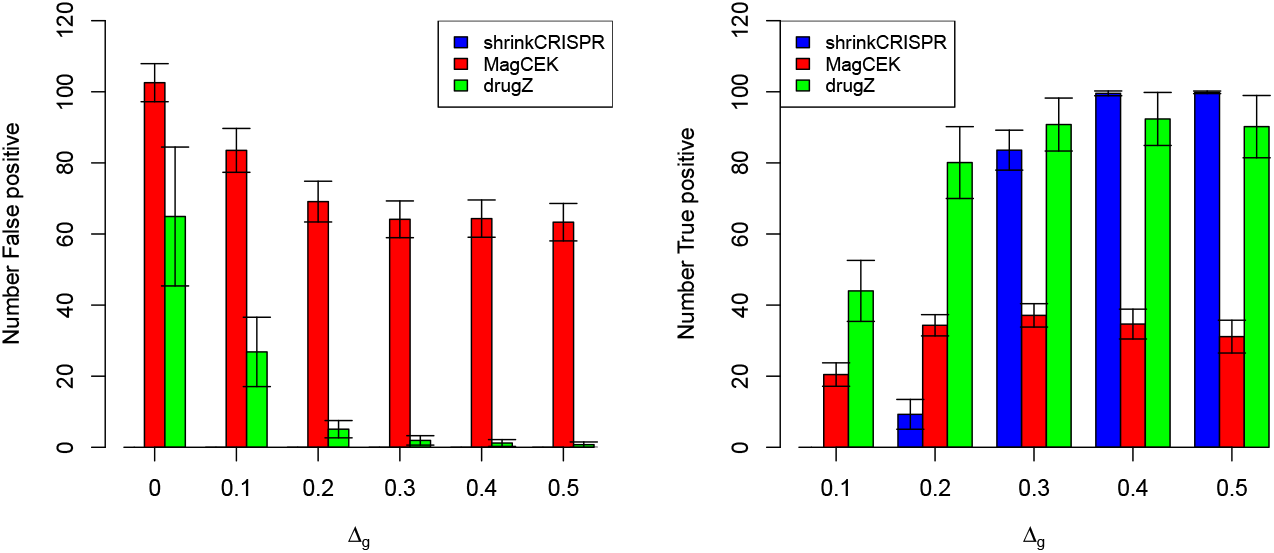
Results per method and simulation scenario, for the paired design. Left panel: average number of false positive hits across 100 simulated datasets. Right panel: average number of true positive hits across 100 simulated datasets. Black lines: standard deviation across simulated dataset.

In terms of true hits, drugZ is the best performing approach to detect small lethality effects, with on average 46.8 and 81.2 true positive hits when Δ_*g*_ =0.1 or 0.2, respectively. However, drugZ is not able to detect all true positive hits when Δ_*g*_ ≥ 0.3, with a maximum around 90.2. ShrinkCRISPR displays low power for Δ_*g*_ ≤ 0.2, similar power to drugZ when Δ_*g*_ = 0.3 and it outperforms all methods when Δ_*g*_ ≥ 0.4. MAGeCK shows in all scenarios a low ability to detect true hits, with less than 40% of genes with effect detected across all effects considered. All results are available in Table S1.

Figure 3 presents the ROC curves obtained by averaging the ROC curves across the 100 simulated datasets. Overall, ShrinkCRISPR performs at least as well as drugZ in terms of sensitivity and specificity. In particular, as the number of true positives increases, the corresponding number of false positives increases quicker with drugZ than with ShrinkCRISPR – this is clearly visible from the results obtained with Δ_*g*_ =0.1. Thus, while ShrinkCRISPR is more conservative, it yields a higher ratio of true positive hits compared to false positives, for low Δ_*g*_ values. Finally, MAGeCK shows worse performance than both drugZ and ShrinkCRISPR in every scenario, in terms of sensitivity, specificity as well as error control level. This may be explained by the fact that it involves two one-sided tests, which inflates the false positive rate.

**Figure 3.**
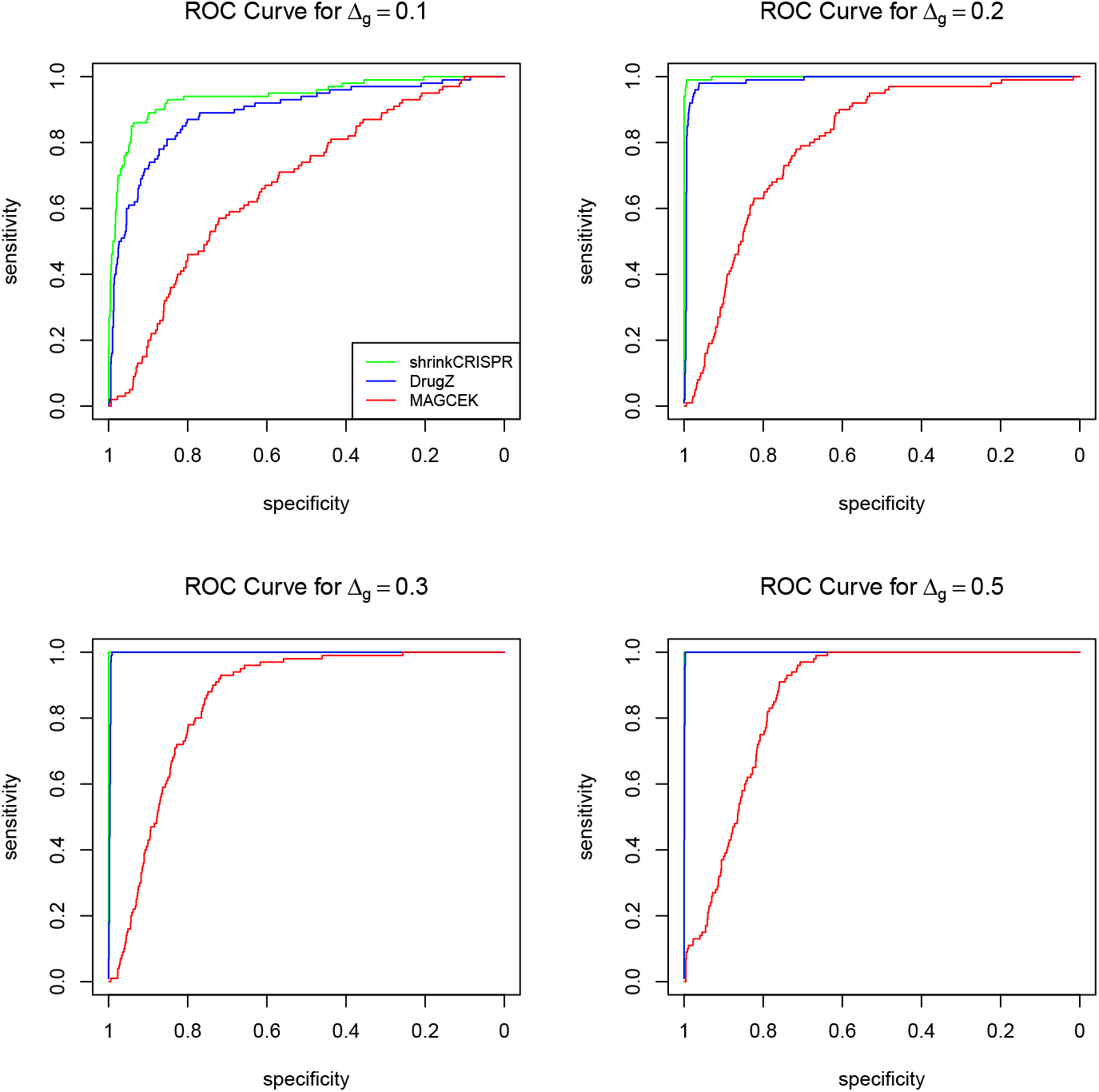
ROC curves per method and simulation scenario with Δ_*g*_ > 0 for the paired design.

We can conclude that drugZ has more power to detect true hits related to small effects (Δ_*g*_ = 0.2), while also yielding more false positives, compared with ShrinkCRISPR in our simulation study with a paired design. For larger effects, ShrinkCRISPR performs better, as it detects as many true hits as drugZ, while yielding virtually no false positives. MAGeCk displays less power and yields a higher false positive rate than the other two methods in the situations considered.

### Independent design

When using an independent design, the initial sgRNA abundance may vary between replicates. Since both MAGeCK and drugZ do not take into account the initial sgRNA abundance, it is to be expected that they perform less well for data produced using this design. Indeed, both MAGeCK and drugZ yield many (> 50 on average, where 50 were expected) false positive hits in simulated datasets (left panel of Figure4). This is understandable as both methods were developed for the paired experimental design. In contrast, ShrinkCRISPR yields no false positives throughout all different simulation scenarios.

**Figure 4.**
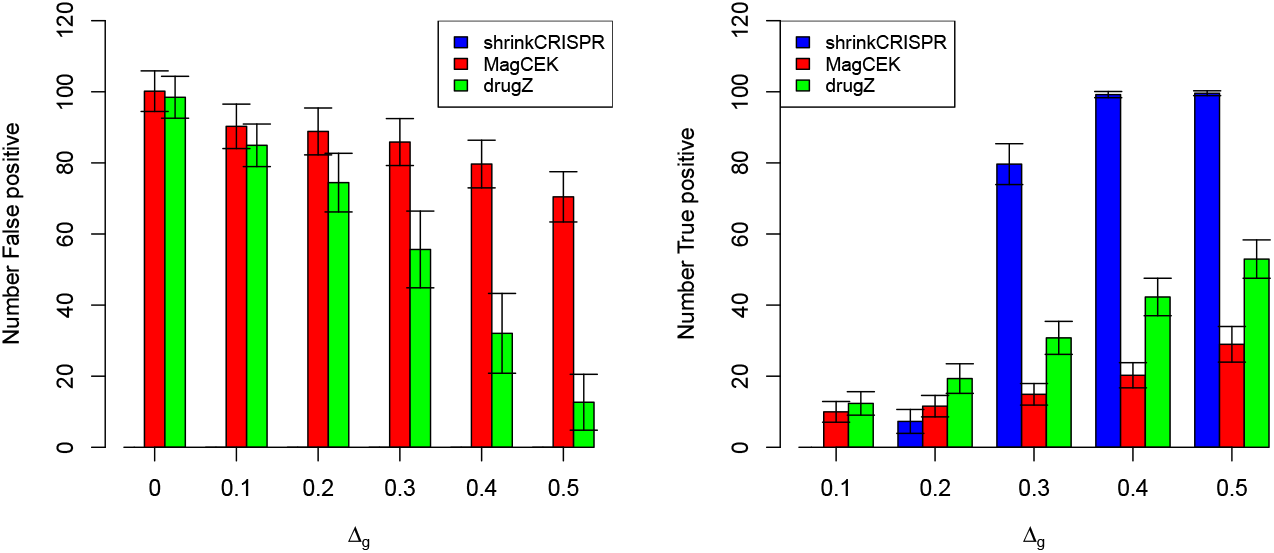
Results per method and simulation scenario, for the independent design. Left panel: average number of false positive hits. Right panel: average number of true positive hits. Black lines: standard deviation across simulated dataset.

In terms of true positives, ShrinkCRISPR is conservative for small effects, detecting at most 5 true hits on average for Δ_*g*_ ≤ 0.2. For Δ_*g*_ ≥ 0.3 and higher, it identifies between 80 and 100% of all hits. MAGeCK and drugZ detect between 10 and 20% of hits for Δ_*g*_ ≤ 0.2, but find only 33.1 % and 55.1 % of all true positives respectively for Δ_*g*_ = 0.5 (Figure 4, right panel).

As expected, for this design ShrinkCRISPR yields ROC curves considerably better compared to both drugZ and MAGeCK (Figure 5). Indeed, ShrinkCRISPR displays similar sensitivity and specificity for the independent design to that for the paired design. In contrast, both drugZ and MAGeCK yield more false positives and less power for the independent design.

**Figure 5.**
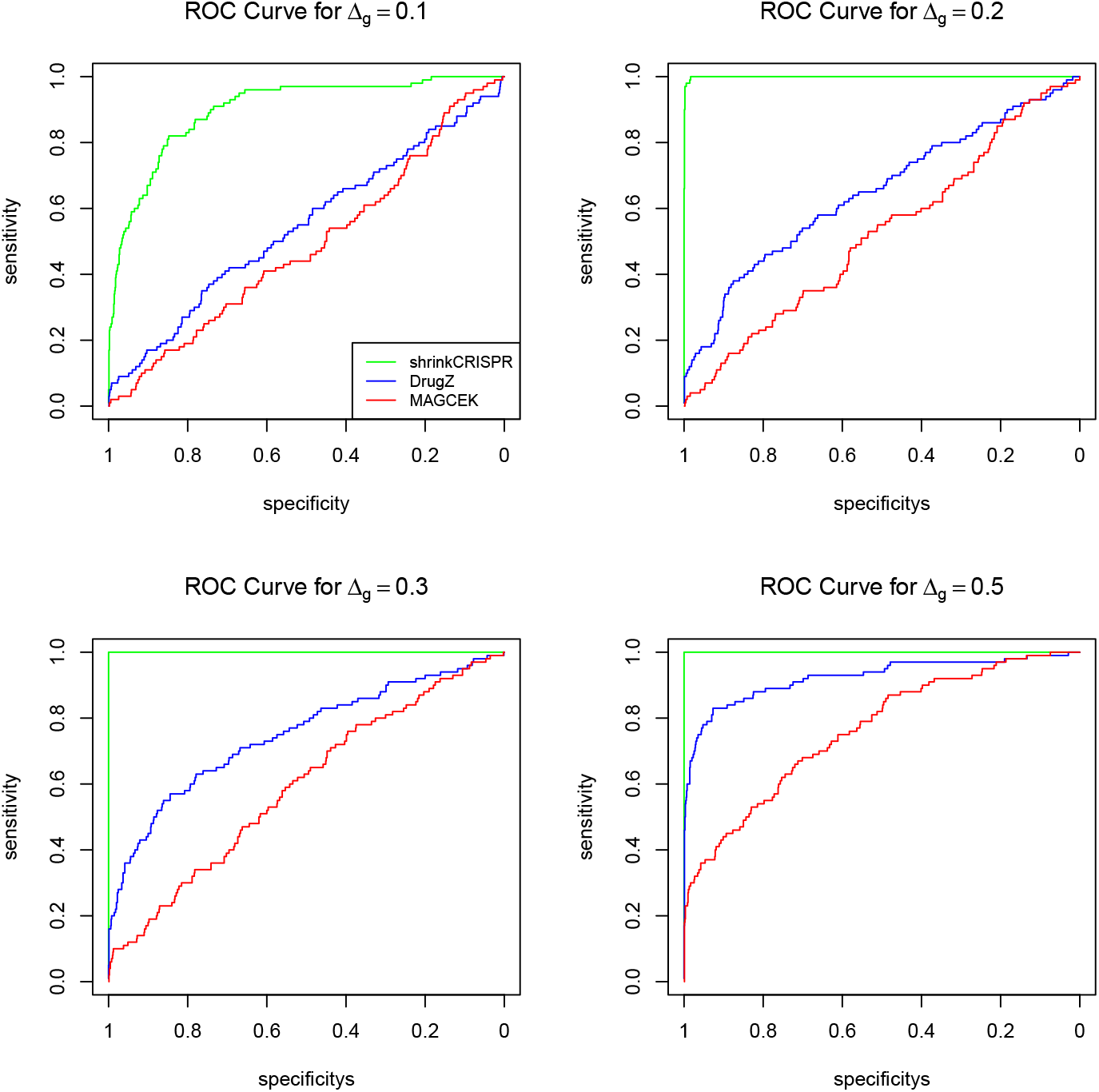
ROC curves per method and simulation scenario with Δ_*g*_ > 0 for the independent design.

Overall, both MAGeCK and drugZ display worse performance with the independent design, compared to the paired design. In terms of false positives, drugZ yields on average 10% false positive hits for Δ_*g*_ = 0, compared to 6.7% in the paired design, and 1% for Δ_*g*_ = 0.5, compared to 0% in the paired design. In addition, the maximum power it achieved in simulations was below 60%. MAGeCK yields more false positives using the independent design, compared with the paired design. Note that the performances of drugZ for an independent design is dependent on the amount of variation at baseline. To illustrate this, we also simulated independent screens with a variance at baseline being approximately half the variance in the results above. This led to an improvement of the performances of drugZ (see Supplementary Table S1).

The performance of both drugZ and MAGeCK is worse under the independent design because both methods do not take into account the initial sgRNA abundance variation. In this case, it is essential to include the initial screen (at *T* = 0) in the analysis.

The absence of false positive hits accross all simulation scenarios highlights the conservatism and potential lack of power of ShrinkCRISPR with a small number of samples. This is confirmed by results obtained using less extreme lethality values for the positive and negative controls (see supplementary table S2). By increasing the variance of lethality effect *sigma*^2^ the number of false positive increases but remains small. It is also important to note that all sgRNAs are simulated with no count increase between T=0 and the later time point T, further limiting the ability to detect hits for shrinkCRISPR as no growth of the different populations of sgRNA transduced cells is simulated.

## Experimental data analysis

We apply ShrinkCRISPR to the following public datasets: 1. cisplatin sensitivity screens, as part of a large study of the sensitivity to a variety of drugs of human RPE1 cells [15]] using a paired design; 2. the longitudinal screens of two cancer cell lines (HeLa and HCT116), performed with the first generation of the Toronto KnockOut (TKO) library [7]. For the latter, we analysed both single time points, as well as multiple time points with the model.

### Cisplatin sensitivity (paired design)

Cisplatin causes DNA crosslinks, of which the repair requires specific DNA damage response genes, most notably components of the Fanconi anemia pathway [9]. This means that, in studies involving screening of cell lines with and without cisplatin, the clearest illustration of biological relevance involves genes whose knockout is specifically lethal in cisplatin-treated cells. In [15] human RPE1 cells were screened using the TKOv3 sgRNA library with and without cisplatin. The different screens were performed in duplicates. As this low number of replicates is challenging for mixed effect models, we grouped two screens that were executed identically (cisplatin 2 and cisplatin 3), yielding four replicates for both treated and untreated cells.

ShrinkCRISPR identified 37 significant hits (lfdr = .10, colored dots in Figure 6), the vast majority showing lower abundance in cisplatin-treated as compared to untreated cells – this is represented by a negative effect appearing on the left side of the volcano plot.

**Figure 6.**
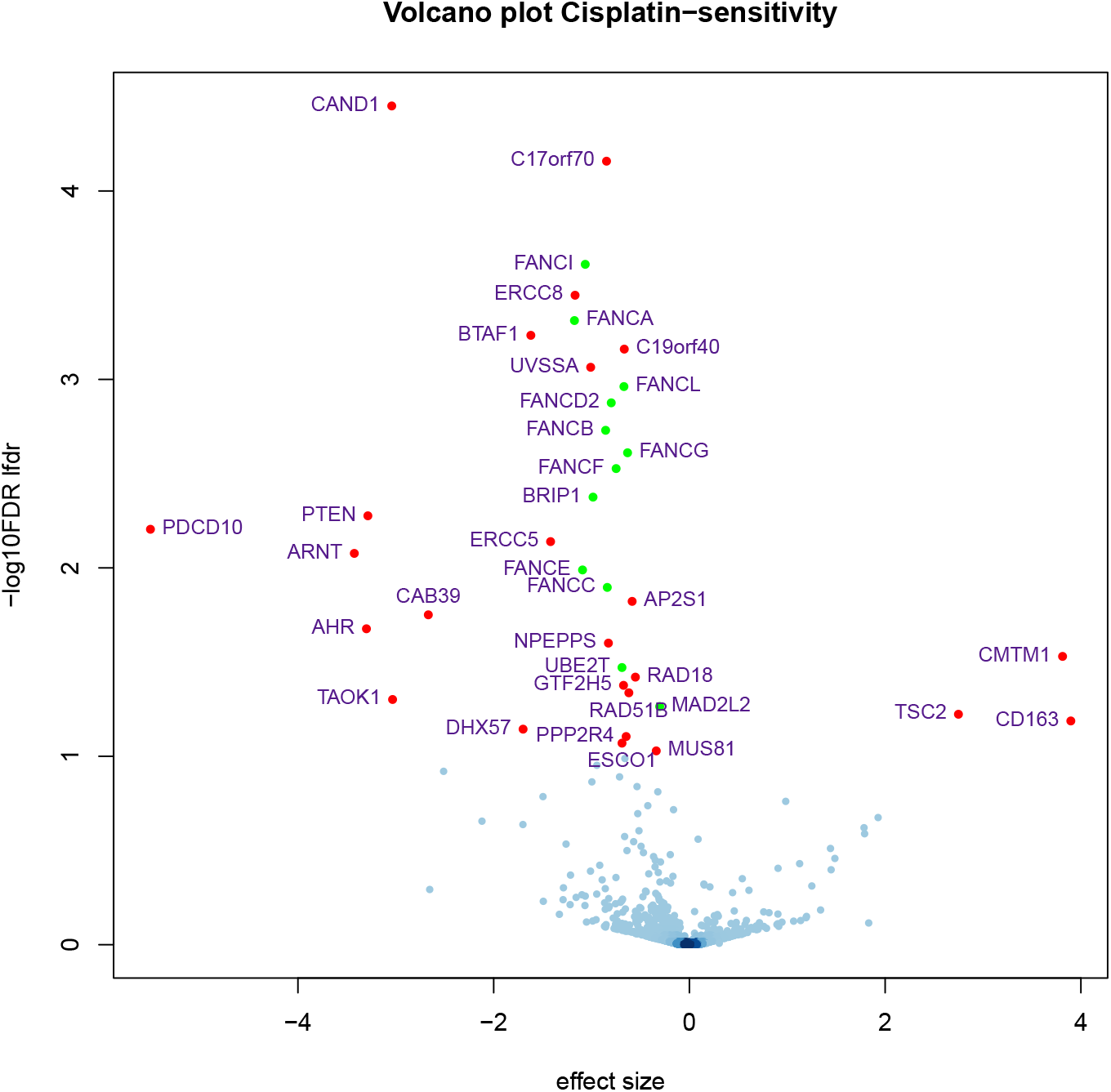
Volcano plot of the analysis of the cisplatin screens, compared to untreated cells. The 37 genes with a lfdr below 0.1 are highlighted; green dots represent Fanconi Anemia genes. Here we point out that *C17ORF70* is now known as Fanconi-Anemia Associated Protein 100 (*FAAP100*)

As expected, these include 12 (out of 22) known Fanconi anemia genes (depicted as green dots in Figure 6), as well as other known DNA repair genes such as *GTF2H5*, *ERCC8*, *C19orf40* (*FAAP24*), *ERCC5*, *RAD18*, *GTF2H5*, *RAD51B* and *MUS81*.

Notably, because the calculated lethality score *Z* is continuous and can take values varying from –∞ to 1 (1 indicating complete depletion relative to controls and to *T*_0_, and negative values indicating enrichment), the effect sizes of enriched genes can potentially be very large. An example of such a hit is *PTEN*, a multifunctional tumor suppressor protein which, when absent, may increase cellular growth and survival (see for example [10]). It displays enrichment in both cisplatin-treated and untreated cells, but this is significantly less pronounced in the treated cells, possibly reflecting that the chemotherapeutic drug cisplatin particularly affects fast growing cells (Figure S4 of the supplementary material).

### Longitudinal analysis (independent design)

To illustrate the use of ShrinkCRISPR with multiple time points, we use screen data produced by [7] with the TKOv1 library. Here we will include data of HCT116 (colorectal carcinoma) and HeLa (cervical carcinoma) cell lines, as both were screened in triplicate at four time points (supplementary table S3). This study used an independent design, with the HeLa cell line used as reference.

The raw data was preprocessed by first calculating fold changes (equation 1) and using rscreenorm [1] to yield quantile-normalized lethality scores. The second step was needed to correct for differences between cell lines. Since the HeLa cell line was used as reference, lethality scores represent the difference in cell counts in the HCT116 compared to HeLa cell line. We first analyzed the different time points individually, yielding four separate sets of results. Subsequently, we used the longitudinal model (equation 4) to analyse all time points together. Genes were selected with lfdr = 0.05.

#### Single time point analysis

The top 10 genes selected per time point display little overlap with those for other time points (Figure 7). In addition, some genes with very small effect sizes are found to be statistically significant. Indeed, *OR52H1* is an olfactory receptor gene, unlikely to be functionally different when knocked out in these cells, and known to be an off-target effect.

**Figure 7.**
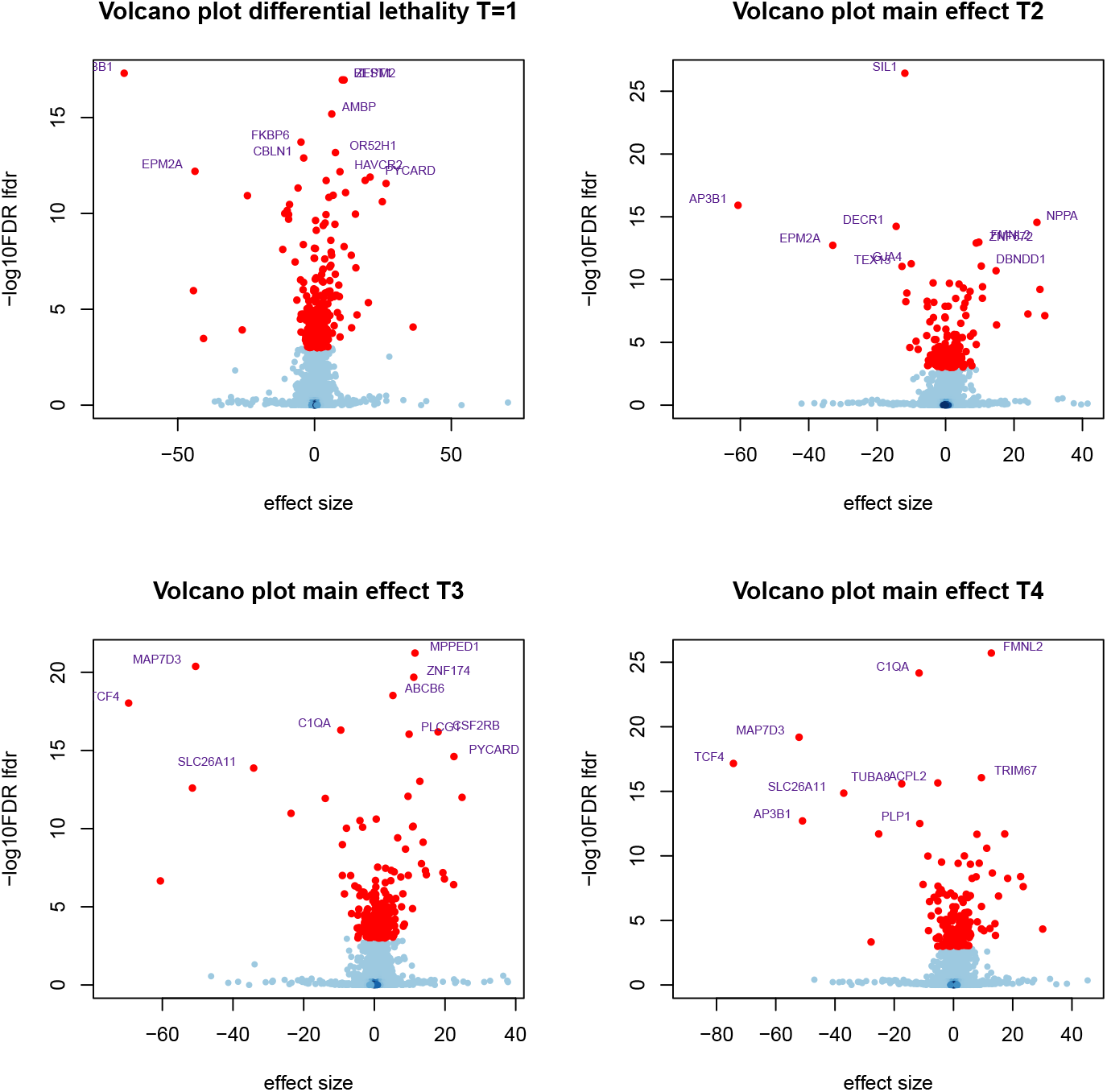
Volcano plots displaying the difference in lethality score between HCT116 and HeLa cells for each time point. Red dots indicate significant genes; the names of the 10 most significant genes are shown. A negative effect size (left-hand side of the volcano) represents either a stronger lethality in HeLa cells, or a growth advantage in HCT116 cells.

Estimated effect sizes for consecutive time points showed remarkable consistency (Figure 8). Indeed, linear regression fitted between estimated effect sizes yielded *R*^2^ = 0.90 between *T* =1 and *T* = 2, *R*^2^ = 0.89 between *T* = 2 and *T* = 3, and *R*^2^ = 0.97 between *T* = 3 and *T* = 4. The high agreement between *T* = 3 and *T* = 4 shows that observed effects mostly occur prior to *T* = 3.

**Figure 8.**
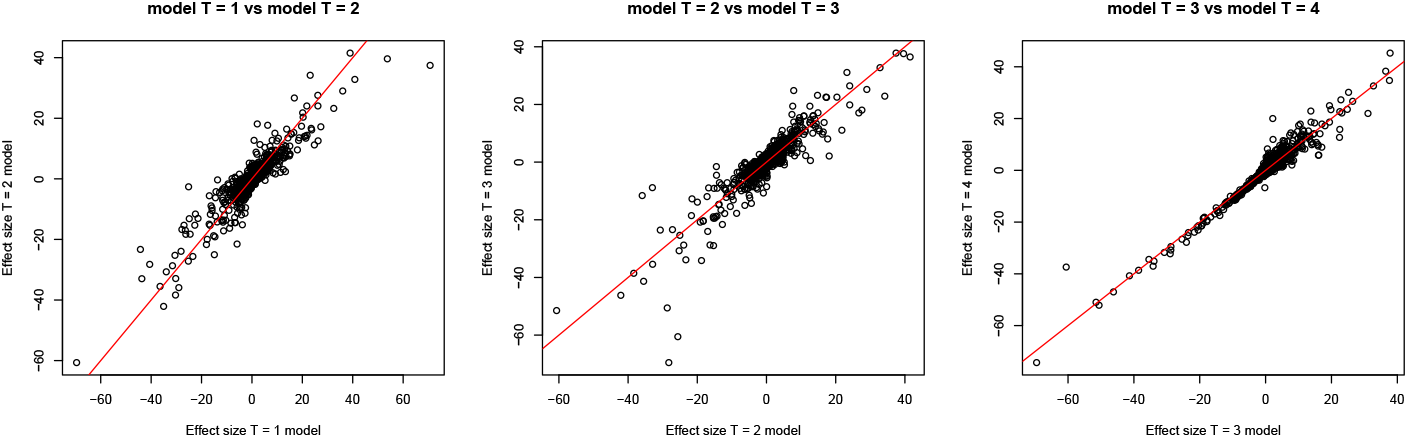
Scatterplots of the effect sizes estimated by ShrinkCRISPR per gene for consecutive time points. The red line represents equal effect sizes.

A comparison between hit lists of genes obtained per time point yielded a substantial number of genes selected only for *T* = 1 (124 out of a total of 336 – table 1). Such differences may reflect early or late biological effects of specific genetic perturbations, such as differences of depletion speeds between cell lines. Numbers of genes selected only for later time points represented smaller proportions of the total of genes selected (28 out of 204, 41 out of 273 and 83 out of 322 for *T* = 2, 3, 4 respectively). Differences between hit lists of genes arise as a result of applying a threshold on the genes’ local FDRs, which takes into account results for all genes at once. Being threshold-dependent, they are less general: indeed, the overlap between gene lists is 100% for a threshold of either 0 or 1. In addition, the lfdr for a gene varies if results for other genes vary. Thus, comparisons between effect sizes as in figure 8 are fairer as they better reflect gene-specific effects. Note also that, while only 36 significant genes are selected by all 4 time points, higher agreement is observed for later time points (Figure S5 of supplementary material).

**Table 1.** Number of significant genes selected by two consecutive time points with lfdr=0.05. Each row displays the number of selected genes for that time point, individually (diagonal) and in overlap with subsequent time points. The total number of significant genes per time point is displayed in the column Significant genes, and the number of significant genes selected by all time points is displayed in the row All.

| Time Point | $T = 1$ | $T = 2$ | $T = 3$ | $T = 4$ | Significant genes |
| --- | --- | --- | --- | --- | --- |
| $T = 1$ | 124 | 138 | 141 | 148 | 336 |
| $T = 2$ | - | 28 | 97 | 97 | 204 |
| $T = 3$ | - | - | 41 | 160 | 273 |
| $T = 4$ | - | - | - | 83 | 322 |
| All | - | - | - | - | 36 |

#### Multiple time points analysis

We selected the most extreme time points to fit the model considering multiple time points. For this part, the rscreenorm quantile normalization step was removed from the preprocessing, and this model (longitudinal) as well as the analysis of single time points *T* = 1 and *T* = 4 were re-run.

Figure 9 illustrates the agreement between the 3 model results. The longitudinal model yielded many more hits: 781 genes compared to 337 and 234 for *T* = 1 and *T* = 4, respectively. Only 101 genes were selected between all model fits. This is mostly due to the disagreement between results using only 1 time point, as only 49 genes from each of the single time point models are not recovered by the longitudinal model. However, the large number of hits that are uniquely found with the longitudinal model suggests it has increased power for hit identification.

**Figure 9.**
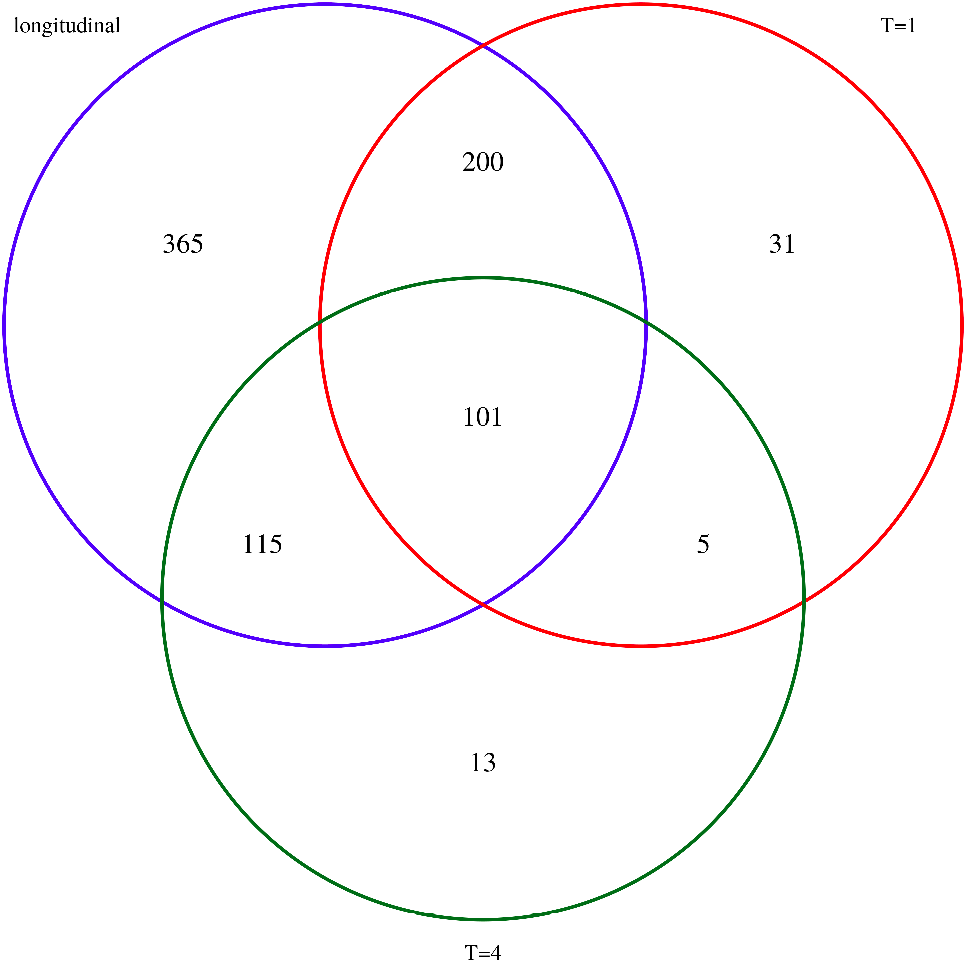
Venn diagram illustrating the overlap of genes found to have a differential effect on cell fitness between HeLa and HCT116 cells, for the three different models (longitudinal, single time point at *T* = 1 and single time point at *T* = 4).

## Discussion

We present ShrinkCRISPR, a new, flexible and powerful method for the analysis of CRISPR screen data for identification of differential effect on cell fitness between conditions. This method incorporates initial sgRNA abundance of each cell line in analyses, enabling its use for various types of experimental designs, including drug-sensitizing screens and isogenic-cell screens. Taking all individual sgRNAs per gene at once in the model, ShrinkCRISPR can test for differences between conditions at the gene level. It makes use of an empirical-Bayes framework, which allows us to represent sgRNA effects as random and condition effects as fixed. The model averages out extreme or conflicting changes, picking out effects that are consistent across most sgRNAs targeting that gene. By adequately accounting for different sources of variability, the method yields as much power as others for most effects, whilst consistently keeping false discoveries under control. Finally, testing at the gene level requires less multiple testing correction than at the sgRNA level, yielding more power.

Our method takes into account existing variation between sgRNAs, as well as possible variation of sgRNAs on cell fitness between conditions via the interaction effect in the model. This yields more robust estimates than those obtained by analysing individual sgRNAs separately: such methods may seemingly produce estimates that display less variability per sgRNA, giving a false impression of more accuracy. In fact, by neglecting inter-sgRNA variability, results represent largely effects on the current experiment, so tend to be difficult to replicate in new experiments, when different replicates, and sometimes different sgRNAs, are used.

In a simulation study, ShrinkCRISPR yielded similar ROC curves to those produced by a another method, drugZ, for drug sensitizing screens using paired designs. However, ShrinkCRISPR yields much less false positives in general. It also outperforms both drugZ and another method, MAGeCK, in the context of independent designs, used e.g. for isogenic screens, as it is the only approach to take into account initial sgRNA abundance. While multiple factors may lead to variability in initial sgRNAs abundance, in published work we found no results reporting such checks. The publicly available data we used in our examples illustrates this point.

The method drugZ was developed to analyse screen data from paired designs. As such, it is not unexpected to perform less well for the analysis of screens generated using independent designs. In our simulation study, we used it to analyse data from independent designs to illustrate the impact of ignoring initial sgRNA abundance on results.

ShrinkCRISPR is the best approach in terms of controlling the proportion of false positive hits, while it is able to find all hits with strong differential effect. However, ShrinkCRISPR is conservative: the false discovery rate is under the desired level, and the method is not able to detect hits with small effect sizes. The low power for detecting small effects could be potentially improved upon by using a spike-and-slab prior for the effect of interest, which would enable the model to better separate a subset of genes with no differential behaviour between groups, from those with differential behaviour. Using the current simulation study setup, however, this did not lead to a better performance (data not shown). The choice of spike-and-slab prior will be available in the R package ShrinkCRISPR.

Results of the simulation study must be interpreted with care. Indeed, each individual simulated dataset used the same effect size Δ_*g*_ for all genes with a condition effect. This enables us to draw conclusions about power of the methods for detecting effects of different sizes, as well as to understand how the amount of false discoveries depends on the effect size present. In practice, experimental data will involve genes with a range of effect sizes. The specific range typically depends on the experimental design and conditions involved. Thus, quantitative results about power and proportion of false discoveries from our simulation study cannot be easily translated to practical applications. The variance existing between sgRNA count within a gene or between replicates can be due to a lot of different phenomenon and is not easily quantifiable. The total amount of variation simulated in the simulation study could not be representative of the experimental results with the improvement of CRISPR screens. However, we have shown with the simulation study that shrinkCRISPR was robust to most variation sources.

Our method has been designed to analyse CRISPR screen data generated by sequencing, consisting of counts. Our proposed pipeline takes the initial sgRNA abundance into account by computing fold changes 1, and subsequently computes lethality scores via rscreenorm. As such, the pre-processed data are no longer counts, and in fact is analysed using model 3 with an error term following a normal distribution.

While we suggest using this pipeline, other researchers may choose to use fewer or none of these pre-processing steps. For example, when studying results for isogenic cell lines, relatively smaller effects are expected than when using cell lines from different individuals or different tissue types. In such cases, sharing of the initial sgRNA abundance eliminates one important variability source. In addition, lethality score distributions for library sgRNAs as well as for assay control sgRNAs tend to be stable across cell lines, and normalization with rscreenorm may be unnecessary. In such cases, the data will involve both over-dispersion as well as potentially zero inflation. The flexibility of the proposed framework enables ShrinkCRISPR to still be used, by fitting model 3 with a negative binomial distribution for the response *Z_gr_* as counts, accounting for over-dispersion. It can also include a term to account for zero inflation.

Some researchers suggest combining multiple test results for sgRNAs targeting the same gene by means of summarizing their p-values (one example is REF), say using the minimum of them. This can lead to over-optimistic results, as the summary works similarly to a filtering of the features, since only one test is selected from a set of them. As a filter, the selection of the sgRNA test with the smallest p-value is not independent of the test result by definition, and this yields a bias on the FDR control method [20].

There are methods currently in use which rely on more sophisticated approaches for combining sgRNA-level results (statistics or p-values) to yield gene-level statistics or p-values [3]. While several methods exist to combine p-values of various tests [6, 14, 16], most of them require independent tests, which is not the case for sgRNAs targeting the same gene. There are p-value combining methods which allow for non-independent test, but then only for one-sided significance testing [2]. Such methods would therefore require two statistical tests, which are clearly not independent. So, using such approaches would increases the severity of multiple testing correction, and possibly lead to an inflation in false positive hits due to correlation between tests.

ShrinkCRISPR relies on enough replicates per combination of group and cell line, ideally 3, to yield reliable results. Indeed, using 2 replicates to a poorer ShrinkCRISPR performance, as variances within and between cell lines are then poorly estimated. In particular, if a single replicate is available for each combination of group and cell line, ShrinkCRISPR cannot be applied. While this can be seen as a too strong requirement by some researchers, we think this is a reasonable restriction: it follows from the need for estimating variability for all sources of variation, which is precisely what enables ShrinkCRISPR to yield less false discoveries. A further challenge when using ShrinkCRISPR is that the effect sizes are not always straightforward to interpret due to the several normalization steps. Furthermore, as all approaches using fold changes, ShrinkCRISPR is sensitive to extreme values for sgRNA initial abundance, in particular very low ones.

The TKO data analysis showed that our approach can account for multiple effects in CRISPR screens, both at the sgRNA and at the replicate levels. Indeed, estimated effects of different time points showed strong agreement: their correlation was at least 90% on average. Finally, by taking multiple time points into account in the model, ShrinkCRISPR significantly increased the power to detect differential effect on cell fitness, finding more time-independent effects than when individual time points were used.

Another important step of pre-processing common to all methodologies based on fold changes is the handling of low counts. Indeed in shrinkCRISPR we create a fold change to measure the population growth of cell transduced with a specific sgRNA. The presence of low sgRNA counts at the initial time point can lead to a large fold change value and thus to an artificially large lethality score. This can then produce false positive hits. By modelling the variance between sgRNAs within a gene, ShrinkCRISPR is more robust to extreme values for individual sgRNAs. However, this may not be sufficient. One common approach to deal with such problems is to add to all raw low counts at *T*_0_ a fixed (small) number of ‘cells’ in order to reduce the impact of such low counts on results.

We conclude that ShrinkCRISPR yields at least as much power to other existing ones for most effects, with much better true positive proportions, even if conservative. As downstream validation studies are extremely time-consuming, it represents an important step towards making better use of data produced, producing more reproducible results, and leading to more efficient studies.

## Author’s contributions

RM had the initial idea for the method, which was further developed by RT. RM and RT wrote scripts, and RT performed analyses. RW, JL and JS provided much needed biological insight. All authors read and approved of the final manuscript.

## Acknowledgements

We are very grateful for the fruitful discussions with Mark van de Wiel on the statistical methodology, and to Klaas de Lint on biological insights. This work was funded by KWF project number 10701.

