## Supplementary material for "ShrinkCRISPR : A flexible method for differential fitness analysis of CRISPR-Cas9 screen data"

In this supplementary materials, we are presenting more detailed simulation results as well as additional information on the experimental data analysis.

#### Contents

**A Simulations Results** **2**

**B Experimental Analysis** **6**

B.1 Cisplatin sensitivity (paired design) . . . . . . . . . . . . . . . . . . . . . . . . . . . . . . . . . 6

B.2 Longitudinal analysis (independent design) . . . . . . . . . . . . . . . . . . . . . . . . . . . . 6

#### A Simulations Results

A simulation study has been set up to compare the performances of our proposed method with the different appraoches MAGeCK and drugZ in various settings. The different approaches were compared in terms of False positives and True positives genes detected in the simulated datasets. The simulations studies are presented in the section 3.1 of the paper. Figure S1 represents the distribution of the simulated lethality scores for the control cell type. Table S1 shows the simulations results for all scenarios. The reduced variance column shows the results obtained when reducing the variance of the sgRNA abundance at baseline between cell lines for an independent design screen. Figure S2 and Figure S3 are an alternative to the bar plots presented in the paper to display the results. Table S2 shows the number of false positive obtained by shrinkCRISPR when the standard deviation of the distibution of fold changes is increased to 0.5 and 1.

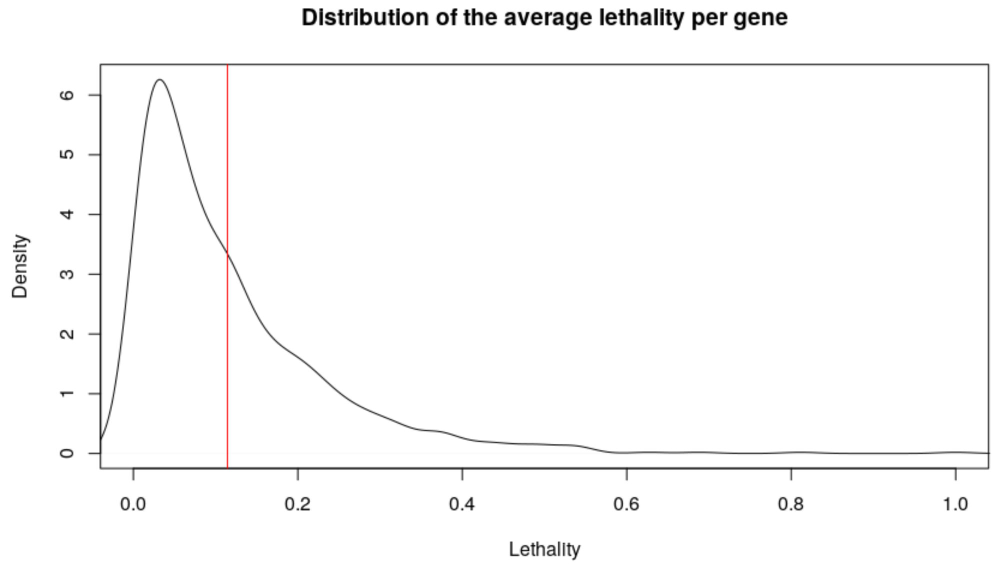


Figure S1: Density plot of the simulated average lethality scores from the 1000 simulated genes in a control cell replicate. The red horizontal line represent the mean of the distribution.

Table S1: Average number of false positive hits (FP) and true positive hits (TP) for each simulation scenario. Into brackets are the standard deviations across datasets for each scenario.

| Method | $\Delta_g$ | Paired Design | | Independent Design | | Reduced variance | |
|---|---|---|---|---|---|---|---|
| | | TP | FP | TP | FP | TP | FP |
| ShrinkCRISPR | 0 | - | 0(0) | - | 0(0) | - | - |
| | 0.1 | 0(0) | 0(0) | 0(0) | 0(0) | - | - |
| | 0.2 | 8.1(4.7) | 0(0) | 7.0(3.9) | 0(0) | - | - |
| | 0.3 | 81.7(6.9) | 0(0) | 78.0(7.4) | 0(0) | 80.1(6.2) | 0(0) |
| | 0.4 | 99.6(0.8) | 0(0) | 99.1(0.6) | 0(0) | - | - |
| | 0.5 | 99.9(0.4) | 0(0) | 99.8(0.5) | 0(0) | - | - |
| MaGCEK | 0 | - | 105.2(18.9) | - | 101.4(6.7) | - | - |
| | 0.1 | 20.7(7.5) | 85.1(8.0) | 11.3(3.7) | 89.7(6.3) | - | - |
| | 0.2 | 33.3(8.9) | 71.4(2.1) | 12.8(4.3) | 89.3(7.6) | - | - |
| | 0.3 | 35.8(7.1) | 67.2(1.3) | 17.1(4.1) | 85.7(12.6) | 32.5(5.0) | 71.4(6.6) |
| | 0.4 | 33.0(6.7) | 65.6(0.4) | 22.7(3.6) | 79.9(12.2) | - | - |
| | 0.5 | 29.6(9.4) | 65.6(0.5) | 33.1(4.9) | 70.9(7.3) | - | - |
| drugZ | 0 | - | 66.9(6.1) | - | 101.3(7.0) | - | - |
| | 0.1 | 46.8(3.3) | 30.5(6.1) | 13.3(3.2) | 86.3(7.0) | - | - |
| | 0.2 | 81.2(2.5) | 6.1(5.9) | 21.8(2.8) | 75.3(7.2) | - | - |
| | 0.3 | 90.2(3.5) | 2.1(7.0) | 31.3(3.4) | 54.1(6.7) | 61.8(3.2) | 12.8(6.6) |
| | 0.4 | 89.9(3.8) | 1.5(5.6) | 42.9(4.5) | 30.2(6.29) | - | - |
| | 0.5 | 89.4(3.7) | 0.7(5.7) | 55.1(5.6) | 11.7(5.9) | - | - |

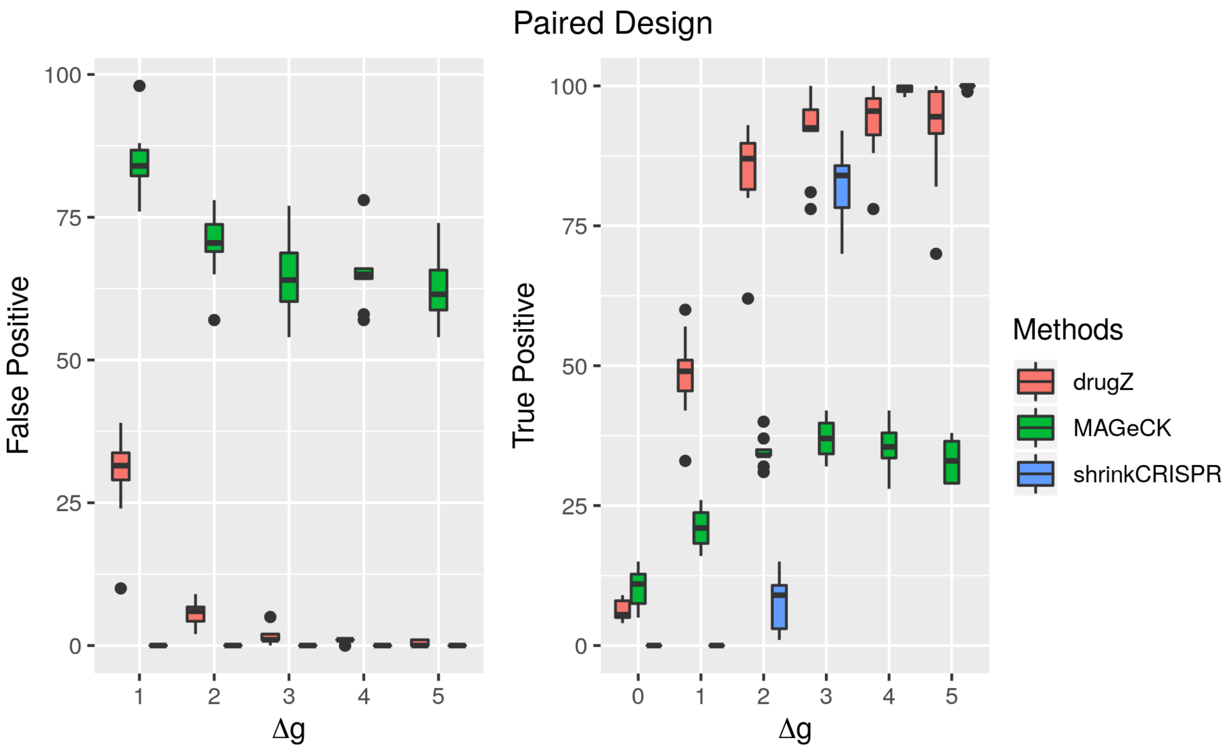


Figure S2: Results per method and simulation scenario, for the paired design. Left panel: average number of false positive hits. Right panel: average number of true positive hits. Standard deviation of the number of true and false positives across simulated datasets are represented by the black lines. 100 datasets were simulated for each scenario.

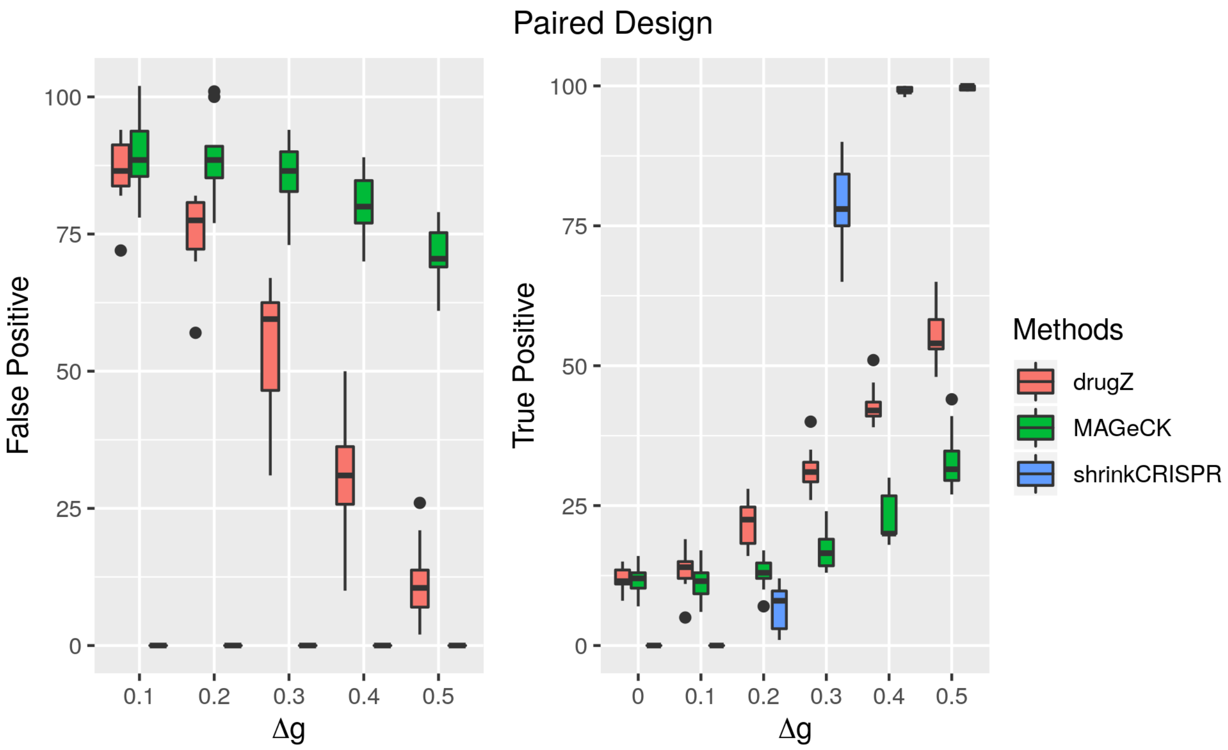


Figure S3: Results per method and simulation scenario, for the paired design. Left panel: average number of false positive hits. Right panel: average number of true positive hits. Standard deviation of the number of true and false positives across simulated datasets are represented by the black lines.

Table S2: Average number of false positive hits (FP) for each simulation scenario with the standard deviation of the fold changes equal to 0.5 and 1. Into brackets are the standard deviations across datasets for each scenario. 10 datasets were simulated for each scenario.

| $\Delta_g$ | $\sigma = 0.5$ | $\sigma = 1$ |
|---|---|---|
| 0 | 2.3(1.7) | 5.9(1.6) |
| 0.1 | 3.0(1.8) | 6.0(1.8) |
| 0.2 | 3.7(2.1) | 6.5(2.3) |
| 0.3 | 4.8(2.3) | 6.6(1.9) |
| 0.4 | 7.6(3.6) | 7.3(2.7) |
| 0.5 | 12.9(3.8) | 8.1(3.3) |

### B Experimental Analysis

#### B.1 Cisplatin sensitivity (paired design)

We performed, using shrinkCRISPR, the analaysis of cisplatin sensitivity screens, as part of a large study of the sensitivity to a variety of drugs of human RPE1 cells (see main manuscript). From this analysis 37 genes were found significant. Figure S4 displays the fitness scores obtained by shrinkCRISPR for the 37 genes in the two cell line conditions : treated with cisplatin and without cisplatin.

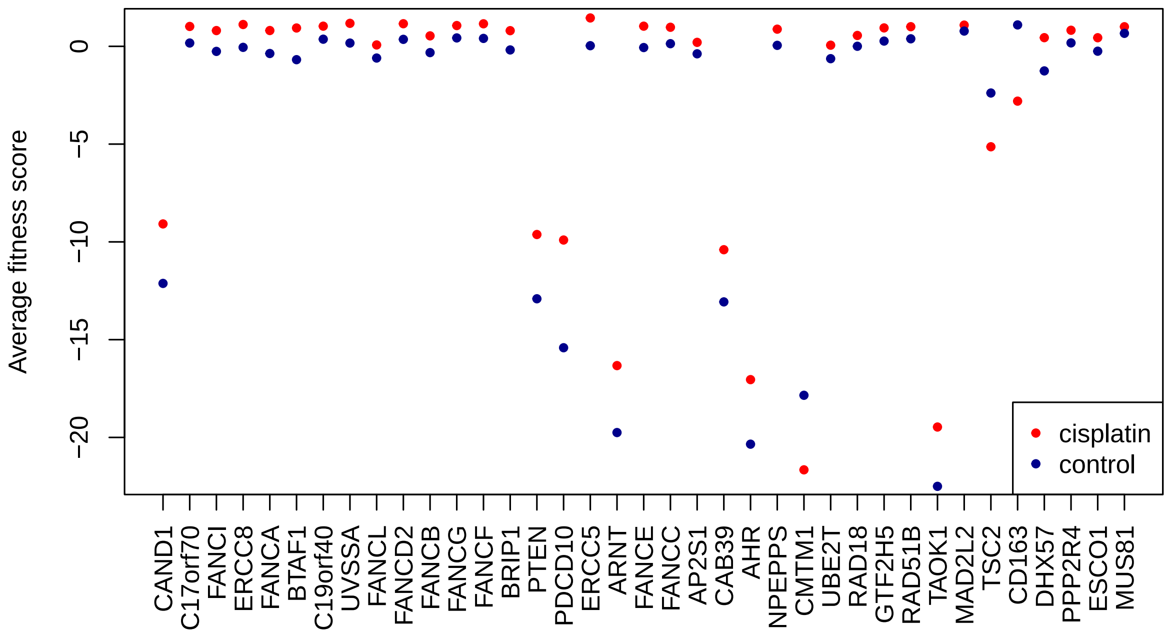


Figure S4: average fitness score for both cisplatin treated cells (red) and control cells (blue). A positive fitness score indicates depletion of the sgRNA over time; negative values indicate enrichment.

#### B.2 Longitudinal analysis (independent design)

We performed the analysis of the publicly available screens of the Toronto KnockOut (TKO) library. 6 differents cell lines (DLD1, GBM, HCT116_1, HCT116_2, HeLa and RPE1 have been screened at several time points. Each cell lines consisted of duplicates or triplicates. A summary of the number of samples available for each cell line at each time points is presented in table S3. Figure S5 illustrate the intersection of the significant genes obtained by shrinkIso for each spearate time points.

Table S3: Number of replicates at each measured time points for each cell line screens in the TKO library publicly available dataset.

| Time Point | DLD1 | GBM | HCT116_1 | **HCT116_2** | **HeLa** | RPE1 |
|---|---|---|---|---|---|---|
| $T = 0$ | 1 | 1 | 1 | **1** | **1** | 1 |
| $T = 1$ | 1 | 2 | 2 | **3** | **3** | 2 |
| $T = 2$ | 1 | 2 | 2 | **3** | **3** | 2 |
| $T = 3$ | 1 | 2 | 2 | **3** | **3** | 2 |
| $T = 4$ | 0 | 0 | 2 | **3** | **3** | 2 |
| $T = 5$ | 0 | 0 | 2 | **0** | **0** | 0 |

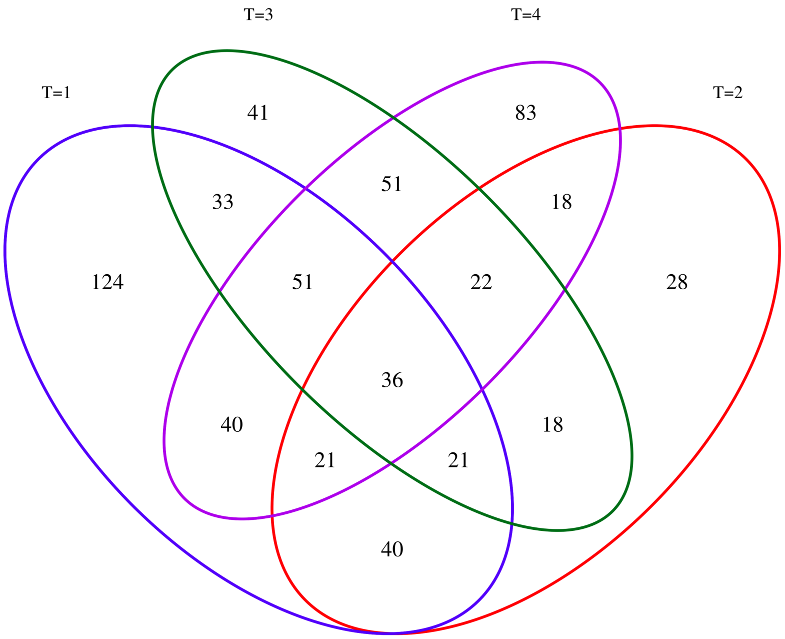


Figure S5: Venn diagram illustrating the overlap between the genes presenting a differential effect on cell fitness between the HeLa and HCT116_2 cell lines for the 4 analyses ran at the 4 different time points.
